# Draft Genome Sequences of Nine Japanese Strains of the Kiwifruit Bacterial Canker Pathogen, *Pseudomonas syringae* pv. actinidiae Biovar 3

**DOI:** 10.1101/2020.08.27.270900

**Authors:** Takashi Fujikawa, Hiroyuki Sawada

**Affiliations:** Institute of Fruit Tree and Tea Science (IFTS), National Agriculture and Food Research Organization (NARO), Tsukuba, Ibaraki, Japan; Genetic Resources Center (GRC), NARO, Tsukuba, Ibaraki, Japan

## Abstract

*Pseudomonas syringae* pv. actinidiae is the pathogen that causes kiwifruit bacterial canker and is categorized into several groups (biovars). In Japan, biovar 3, known as the pandemic group, was first discovered in 2014. Here, we sequenced the genomes of nine Japanese biovar 3 strains.

---

The kiwifruit bacterial canker pathogen, *Pseudomonas syringae* pv. actinidiae, causes serious damage to kiwifruit production worldwide (1), and is currently subdivided into several groups (biovars) (2, 3). Among them, biovar 3 caused recent pandemics of this disease in various parts of the world (1–5). In Japan, biovar 3 strains were first discovered in 2014 (6) and have caused enormous damage since then (3). Because biovar 3 strains were not detected at all in Japan until 2014 (3, 6) and it had been clarified that the pandemic lineage of biovar 3 originated in China (4, 5), it is speculated that biovar 3 might have invaded Japan from any country where it previously occurred (3). In biovar 3, various types of the integrative and conjugative element (ICE) with differing structures and insertion sites have been detected (5). Among them, Pac_ICE1 was detected by PCR assays in biovar 3 strains isolated in Japan (6). Pac_ICE1 has also been detected in biovar 3 strains isolated in China and New Zealand (5). Here, we selected nine Japanese strains of biovar 3 (Table 1) from the NARO Genebank collection (https://www.gene.affrc.go.jp/index_en.php) and sequenced their genomes to further advance our understanding of biovar 3 genomics.

**Table 1.** Genome data and accession numbers of nine strains of *Pseudomonas syringae* pv. actinidiae biovar 3.

| Strain information |  |  | Genome information |  |  |  |  | PGAP<sup>a</sup> annotation |  |  | Reads information |  |  |  |
| --- | --- | --- | --- | --- | --- | --- | --- | --- | --- | --- | --- | --- | --- | --- |
| Strain | Isolation host | Isolation area and year | GenBank accession no. | Genome size (bp) | G+C content (mol% ) | No. of contigs | *N*<sub>50</sub> | Total no. of genes | rRNAs (5S, 16S, 23S) | tRNAs | SRA<sup>b</sup> accession no. | No. of reads | Average length (bp) | Genome coverage (×) |
| MAFF 212101 | *Actinidia chinensis* | Saga Prefecture (Pref.), 2014 | PGSP00000000 | 6,112,610 | 58.6 | 461 | 24,702 | 5,931 | 4, 1, 1 | 38 | SRR11744872 | 2,256,006 | 152.0 | 54.7 |
| MAFF 212104 | *A.chinensis* | Ehime Pref., 2014 | PGSX00000000 | 6,135,799 | 58.6 | 489 | 23,513 | 6,021 | 2, 1, 1 | 46 | SRR11744878 | 4,126,079 | 241.2 | 159.9 |
| MAFF 212109 | *A.chinensis* | Wakayama Pref., 2014 | PGSS00000000 | 6,142,013 | 58.6 | 457 | 25,821 | 6,081 | 3, 1, 2 | 48 | SRR11744873 | 6,152,457 | 280.9 | 278.1 |
| MAFF 212111 | *A.chinensis* | Fukuoka Pref., 2014 | PGSQ00000000 | 6,134,312 | 58.5 | 428 | 29,979 | 6,019 | 3, 1, 1 | 40 | SRR11744874 | 4,591,637 | 269.7 | 198.8 |
| MAFF 212115 | *A.chinensis* | Fukuoka Pref., 2014 | PGSO00000000 | 5,786,754 | 58.7 | 495 | 22,601 | 5,662 | 2, 1, 1 | 37 | SRR11744870 | 1,763,341 | 181.5 | 54.0 |
| MAFF<br>212118 | *A.chinensis* | Fukuoka<br>Pref.,<br>2014 | PHQZ000<br>00000 | 4,223,244 | 58.6 | 751 | 12,356 | 5,847 | 1, 1, 2 | 37 | SRR117<br>44871 | 1,002,668 | 172.0 | 52.5 |
| MAFF<br>212145<br>(Saga2-1) | *A.chinensis* | Saga Pref.,<br>2014 | PGSV0000<br>0000 | 6,052,839 | 58.5 | 632 | 16,127 | 5,980 | 2, 1, 2 | 38 | SRR117<br>44876 | 3,753,601 | 172.2 | 38.8 |
| MAFF<br>212357<br>(1404) | *A.deliciosa* | Shizuoka<br>Pref.,<br>2014 | PGSZ0000<br>0000 | 6,127,733 | 58.6 | 408 | 28,264 | 5,981 | 3, 2, 1 | 52 | SRR117<br>44880 | 3,200,751 | 187.1 | 96.4 |
| MAFF<br>212440<br>(psa14202<br>7) | *A.chinensis* | Ehime<br>Pref.,<br>2014 | PGSW000<br>00000 | 6,130.793 | 58.5 | 481 | 23,234 | 6,018 | 4, 2, 1 | 54 | SRR117<br>44879 | 2,432,513 | 223.1 | 85.8 |

<sup>b</sup>Sequence Read Archive

Genomic DNAs of the nine strains were prepared and sequenced following the methods of our previous study (7). Briefly, the strains were cultivated in YP broth at 27 ◻ for 1 day with agitation at 140 rpm. Then, 1-ml aliquots of each culture were used for genomic DNA extraction with a DNeasy mini kit (Qiagen, Hilden, Germany). The DNA library was prepared from genomic DNA using an Ion Plus Fragment Library Kit and sequenced using an Ion PGM sequencer with an Ion PGM Hi-Q View OT2 kit, an Ion PGM Hi-Q View Sequencing kit, and a 318 Chip kit v2 (all from Thermo Fisher Scientific Inc., Waltham, MA, USA), according to the manufacturer’s instructions. The sequence reads were quality controlled (a quality score < 20) and adapter sequences were removed using CLC Genomics Workbench version 12 (Qiagen). Using these reads, multiple contigs were assembled *de novo* using the same software with default parameters (mapping mode = create simple contig sequences (fast), automatic bubble size = yes, minimum contig length = 500, automatic word size = yes, performing scaffolding = yes, auto-detect paired distances = yes). The draft genomes were annotated using the NCBI Prokaryotic Genome Annotation Pipeline (PGAP) v4. 1 (8). The guanine and cytosine (G+C) content and genome size for these strains were found to be 58.5–58.7% and 4.2–6.1 Mbp, respectively (Table 1). PGAP identified 5,662–6,081 genes, including multiple rRNA and tRNA genes. Various polymorphisms were detected in the Pac_ICE1 regions of these strains, except MAFF 212115 and 212118, from which Pac_ICE1 could not be detected. Further investigations are needed to determine whether MAFF 212115 and 212118 possess Pac_ICE1. Other than Pac_ICE1, no ICEs were detected in the nine draft genomes sequenced in this study. This information will contribute to future studies on the origin, evolution, transmission, and pathogenicity of biovar 3 worldwide.

## Data availability

All sequences identified in this study have been deposited in GenBank (see Table 1 for accession numbers).

## Acknowledgments

We are grateful to Ms. H. Hatomi and Ms. A. Sasaki for supporting this work. We also thank the members of IFTS-NARO and GRC-NARO for their helpful discussions. We would like to thank Editage (www.editage.jp) for English language editing. This research received no specific grant from any funding agency in the public, commercial, or not-for-profit sectors.

